# Flow-driven lumen remodeling and valve opening in the vas deferens

**DOI:** 10.64898/2026.03.23.713590

**Authors:** Nicole Ng Shu Ying, Qin Ying Lim, Gen Yamada, Tsuyoshi Hirashima

## Abstract

Biological ducts must transport fluids while preserving structural integrity, yet how mechano-signaling coordinates wall deformation with luminal flow in vivo remains unclear. Here we combine intravital two-photon excitation microscopy, light-sheet imaging and FRET-based kinase biosensors to resolve ejaculation-like events in the mouse vas deferens. Acute phenylephrine stimulation elicits a sequence of luminal dynamics: an initial retrograde pressure-redistribution wave followed by a ballistic antegrade flow that propels dense sperm suspensions from proximal to distal duct. This contraction-driven flow opens a normally collapsed, wrinkled distal segment, driving progressive lumen expansion and unfolding of epithelial wrinkles. We show that the vas deferens actively modulates luminal geometry in response to these flow dynamics: ROCK activity in smooth muscle is required for global contraction and cAMP-associated signaling modulates this contractile response. By contrast, ERK activity in circumferential smooth muscle is dispensable for the ductal contraction but essential for active, flow-dependent remodeling of the distal lumen, forming the core of the mechano-signaling module that couples sperm flow to valve opening. These findings establish the vas deferens as an experimentally tractable model of ductal tissue hydraulics and reveal a mechano-signaling framework by which a tubular organ converts transient muscular input into robust, directional luminal transport.

**SIGNIFICANCE STATEMENT:** Male reproductive ducts must rapidly propel sperm-containing fluids forward, yet how they do so in living animals has remained unclear due to a lack of imaging studies. By combining real-time in vivo imaging and molecular activity reporters, we observe the mouse vas deferens at work and link each phase of ejaculation-like transport to specific signaling pathways. We find that ROCK activity is closely linked to the overall squeeze of the duct, cAMP-associated signaling modulates this contractile response, and ERK is uniquely required to open and remodel a normally closed distal valve in response to flow. This mechano-signaling framework offers a general blueprint for how tubular organs coordinate muscle contraction, tissue shape change, and directional luminal transport.

## INTRODUCTION

Biological ducts and tubular organs, such as the gastrointestinal, cardiovascular, and reproductive tracts, dynamically regulate fluid and cellular transport while preserving structural integrity. Their physiological function relies on a tight coupling between luminal flow, tissue mechanoresponse, and active remodeling of the wall, a regime that can be understood within the broader concept of mechanobiology, especially tissue hydraulics (1–3). In this framework, composite tissue structures consisting of smooth muscle, stroma, and epithelium operate as an integrated mechanical system: contractile forces and pressure gradients generate flow; flow imposes shear and strain on luminal tissue surfaces; and these mechanical cues, in turn, activate signaling pathways that reshape duct morphology (4, 5). Despite its conceptual importance, direct in vivo visualization of how mechano-signaling and tissue mechanics are coordinated spatio-temporally within a single ductal organ remains limited.

The vas deferens offers a powerful model to study duct function on fast timescales. Its native role is to rapidly propel dense suspensions of spermatozoa or sperm cells during ejaculation through regulated smooth muscle contractions (6–8). Building on an earlier hypothesis that coordinated contractions along the duct generate efficient forward transport of sperm cells (9– 11), subsequent biomechanical studies using tracer injections revealed complex patterns of luminal movement, including transient retrograde flows and pressure redistribution (12, 13), providing experimental support and refinement of this model. These works have provided a rich biomechanical description of vas deferens function over the past several decades. However, with the rise of molecular biology and genetics, attention shifted toward identifying channels, receptors, and regulatory pathways (14–16). As a result, our current understanding of the vas deferens remains fragmentary: we know much about the molecular components and the global mechanics, but far less about how they are integrated in vivo to control luminal transport and rapid tissue remodeling within seconds to minutes.

Previous pharmacological and electrophysiological studies have been particularly informative in defining the contractile properties of vas deferens. Systemic or local adrenergic stimulation, combined with tension recordings and nerve-stimulation paradigms, established the biphasic twitch–tonic nature of vas deferens contraction and clarified the cellular and molecular mechanisms that initiate and regulate this contraction (15, 17–19). Yet, these approaches typically report either global force or membrane potential, providing limited information about how luminal sperm cells, epithelial architecture, and distinct smooth muscle layers respond simultaneously under near-physiological conditions. Moreover, anatomical studies have reported that the vas deferens exhibits a highly folded, wrinkled epithelium with an apparently collapsed lumen in both rodents and humans (7, 20, 21). However, how luminal sperm flow is routed through this complex structure, and how signaling networks in the duct wall support rapid luminal sperm transport for ejaculation, remain open questions.

Here, we address these gaps by combining intravital two-photon excitation microscopy (TPEM), light-sheet imaging of optically cleared whole tissues, and FRET-based kinase measurement with pharmacological perturbations in the mouse vas deferens. Using phenylephrine (PE) application to vas deferens as a temporally precise, imaging-compatible trigger of contraction, we resolve the sequence of ejaculation-like transport, from retrograde luminal flow detected within three sec after stimulation and subsequent ballistic antegrade sperm transport to distal lumen opening and epithelial wrinkle unfolding. By monitoring and perturbing several signaling activities in distinct smooth muscle layers, we link specific kinase modules to longitudinal contraction and flow-dependent distal remodeling.

## RESULTS

### Phenylephrine (PE) induces vas deferens contraction and drives luminal sperm transport

We employed PE treatment to elicit vas deferens contraction in vivo in mice. Compared with alternative approaches, such as natural mating–induced ejaculation or electrical stimulation of the sympathetic pathways (22, 23), this method offers a technically simple and temporally precise means to trigger tissue contractility while remaining fully compatible with high-resolution live imaging (24–26). In anesthetized mice, the vas deferens was surgically exposed through a scrotal incision, and 10 µL of phenylephrine solution was applied uniformly onto the exposed tissue surface. (**Figure 1A**). Based on empirical dose testing under the in vivo imaging condition, 10 mM PE was used because lower concentrations up to 1 mM did not elicit a robust and reproducible contraction (**Figure S1A**).

**Figure 1.**
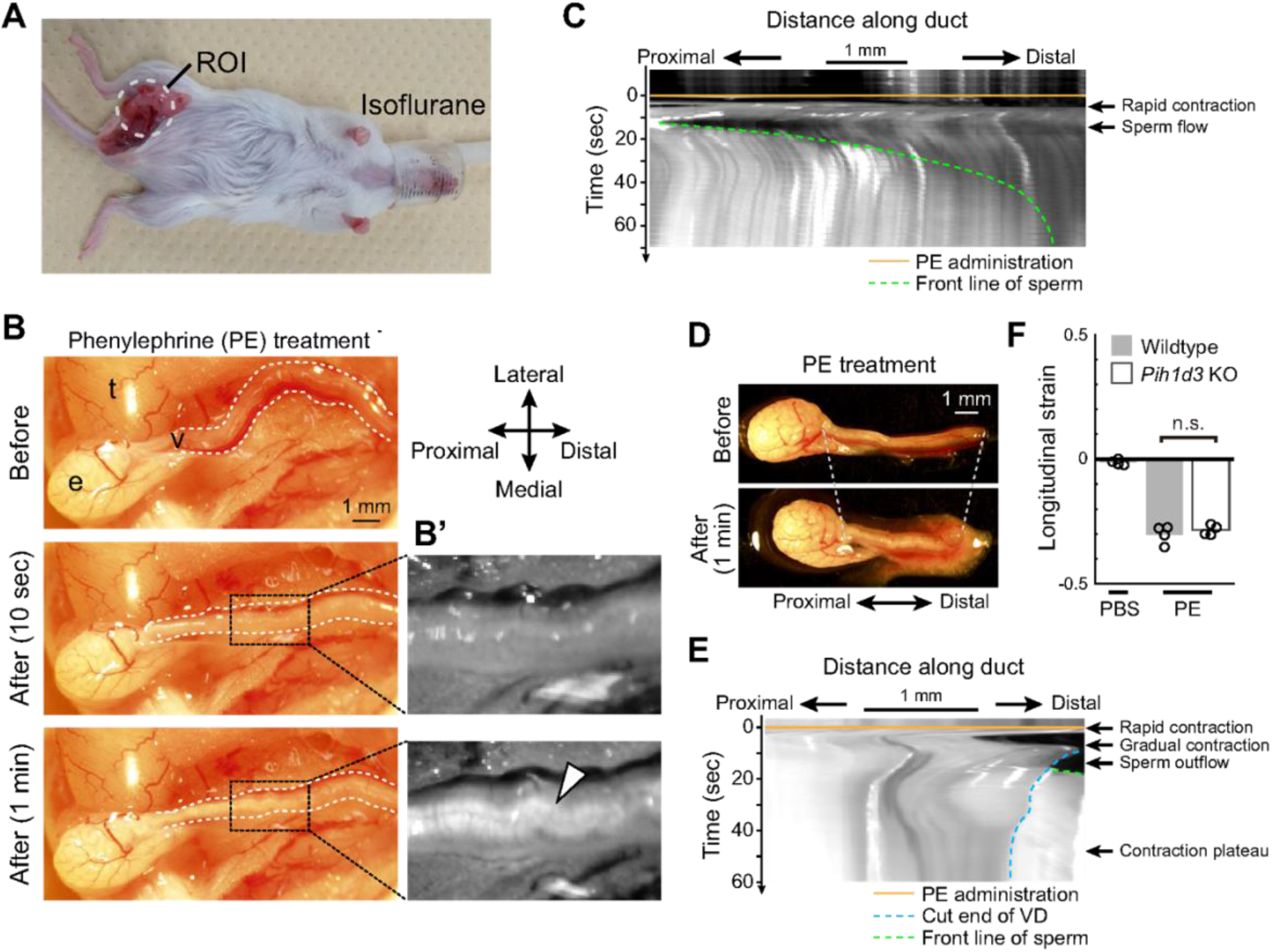
Vas deferens contraction and luminal sperm transport induced by Phenylephrine. (A) Top view of an isoflurane-anesthetized mouse. The region of interest (ROI) highlights the vas deferens within the male reproductive tract. (B) Morphological changes in the vas deferens before and after PE treatment in vivo. e, cauda epididymis; t, testis; v, vas deferens. White dotted lines outline the vas deferens. Scale bar, 1 mm. (B’) Grayscale images showing luminal sperm fluid (arrowhead) entering the mid-vas deferens (region indicated by the black dotted box in B) and flowing from the proximal toward the distal end. (C) Kymograph of vas deferens dynamics during PE treatment in vivo. The orange line marks the timing of PE treatment; the green dotted line indicates the advancing front of luminal sperm fluid. Horizontal scale bar, 1 mm. (D) Morphological changes in an isolated vas deferens preparation before and after PE treatment. White dotted lines mark the proximal end and the distal cut end, respectively. Scale bar, 1 mm. (E) Kymograph of isolated vas deferens during PE treatment ex vivo. The orange line marks the time of PE treatment; the blue dotted line indicates the distal cut end; the green dotted line indicates the front of sperm efflux from the cut end. Horizontal scale bar, 1 mm. (F) Relative longitudinal length of the vas deferens after versus before treatment. PBS served as the vehicle control for wildtype tissues. PE was applied to wildtype and *Pih1d3* knockout vas deferens. Barplots indicate mean value. N=4. Mann-Whitney U-test, p=0.343.

PE treatment induced a rapid and robust contractile response of the vas deferens: the normally undulating duct straightened within five sec of PE treatment (**Figures 1B,B’**). Visual inspection confirmed that a contraction began in the proximal vas deferens and propagated distally along the duct (**Movie 1**). This contraction wave immediately propelled luminal sperm cells from the proximal to the distal vas deferens (**Figures 1B,B’**). Luminal sperm transport became clearly detectable once the contraction stabilized, approximately 10 sec after PE treatment, and persisted for up to 60 sec (**Figure 1C**). Because sperm in the proximal vas deferens can exhibit intrinsic motility, we then asked whether the PE-induced luminal transport reflected sperm self-propulsion or tissue-driven flow. To distinguish between these possibilities, we examined *Pih1d3* homozygous knockout males, which produce immotile sperm (27). An equivalent transport pattern was observed in *Pih1d3* knockout males (**Movie 2**), indicating that PE-induced luminal sperm transport is driven primarily by vas deferens contraction rather than sperm motility.

A similar response was observed in isolated vas deferens preparations retaining the cauda epididymis (**Figure 1D**). For ex vivo assays, 5 µL of 10 µM PE solution was applied uniformly to the isolated tissue. This concentration was selected based on empirical dose testing, which showed that 10 µM PE produced robust and reproducible longitudinal contraction while avoiding unnecessarily high concentrations (**Figure S1B**). PE immediately induced a rapid contraction, followed by a progressive shortening of the longitudinal axis that drove sperm efflux from the cut distal end (**Figures 1D,E; Movie 3**). This mechanically driven outflow was also observed in *Pih1d3* knockout vas deferens (**Movie 4**). PE treatment led to a ~30% reduction in the longitudinal length of the vas deferens, with wild-type and knockout tissues exhibiting the same degree of contraction relative to PBS-treated controls (**Figure 1F**). Together, these data demonstrate that PE robustly induces vas deferens contraction and ejaculation-like luminal sperm transport, yielding a time-controllable and imaging-compatible system to study the dynamics of sperm flow in vivo and ex vivo.

### Intravital imaging reveals dynamic luminal sperm flow during vas deferens contraction

To visualize luminal sperm transport during PE-induced vas deferens contraction, we performed intravital TPEM in Pax2-LynVenus reporter mice, which label the plasma membrane of sperm as well as epithelial cells throughout the male reproductive tract (28, 29) (**Figure 2A**). This imaging approach enables volumetric visualization from the tissue surface to deeper layers, allowing us to observe the longitudinal and circumferential smooth muscle layers, epithelial cells, and luminal sperm cells at single-cell resolution (**Figure 2A’**).

**Figure 2.**
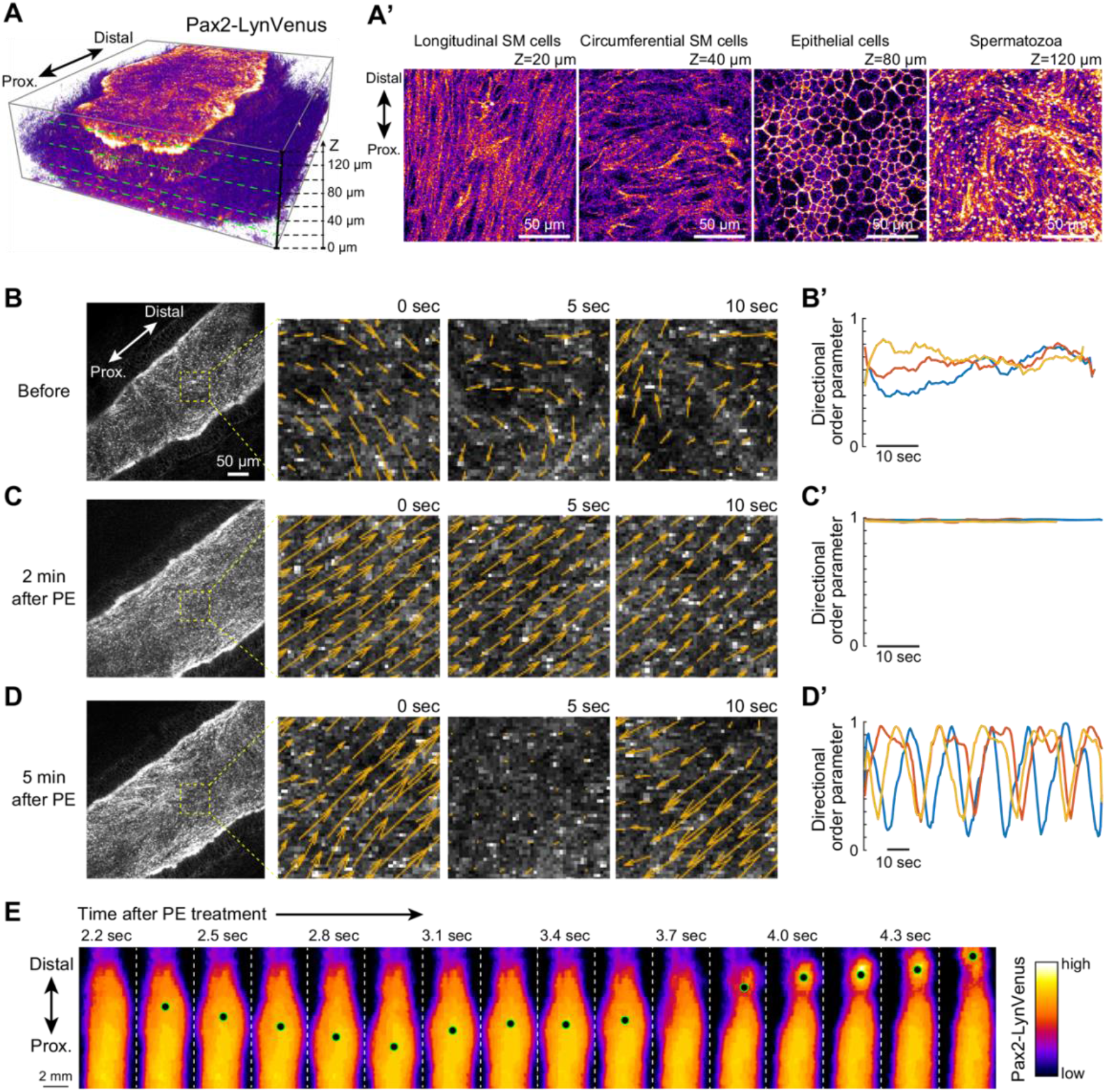
Dynamic luminal sperm fluid driven by vas deferens contraction. (A) Three-dimensional reconstruction of the vas deferens in a Pax2-LynVenus mouse. (A’) Optical sections at increasing depths (Z) from the tissue surface (Z=0 µm), showing the longitudinal smooth muscle (SM) layer at Z=20 µm, the circumferential SM layer at Z=40 µm, epithelial cells at Z=80 µm, and luminal sperm at Z=120 µm. Scale bars, 50 µm. (B–D) Intravital TPEM imaging of luminal sperm dynamics before (B) and after PE treatment (C,D). Ballistic luminal flow immediately after PE treatment occurred too rapidly to be captured; therefore, images are shown from around two min after PE treatment. Orange arrows represent velocity vectors. Scale bar, 50 µm. (B’–D’) Directional order parameter calculated from velocity vectors corresponding to panels (B–D), ranging from 0 (completely disordered motion) to 1 (perfectly aligned collective motion). See Methods for the definition of the directional order parameter. Lines represent individual samples: N=3. Horizontal time scale bars, 10 sec. (E) Macroscopic luminal sperm flow in the vas deferens. Time stamps indicate the elapsed time after PE treatment. Green–black dots mark reference points used to track luminal flow. Color corresponds to Pax2-LynVenus fluorescence intensity. Scale bar, 2 mm.

In the proximal vas deferens, intravital TPEM imaging revealed that luminal sperm move collectively in multiple directions under basal conditions (**Figures 2B,B’; Movie 5**). Because mouse sperm cells become motile during epididymal transit (8), this multidirectional movement reflects intrinsic sperm motility. Upon PE treatment, however, the luminal sperm cells transitioned to a rapid, ballistic flow directed from the proximal toward the distal vas deferens (**Figures 2C,C’; Movie 6**). With the TPEM setup, the tissue contraction immediately following PE delivery caused substantial out-of-plane motion, making it difficult to capture the earliest response within the first 10 sec after stimulation. By five minutes after PE treatment, the ballistic movement switched to an oscillatory, back-and-forth displacement with a mean oscillatory period of 23.4 ± 3.8 sec (mean ± standard deviation) (**Figures 2D,D’; Movie 7**). During this phase, luminal sperm fluid exhibited rhythmic motion driven by periodic smooth-muscle contractions, although their net position over time remained limited. This biphasic response, consisting of an initial rapid twitch contraction followed by a slower, longer-lasting tonic periodic contractions, is consistent with previous descriptions of vas deferens behavior (18, 19).

To capture the earliest response that could not be achieved by TPEM, we utilized high-speed stereomicroscopy. Within three sec of PE treatment, we detected a retrograde luminal flow, directed from the distal toward the proximal vas deferens, followed by a switch to antegrade proximal-to-distal transport (**Figure 2E; Movie 8**). This transient retrograde flow is consistent with previous observations, in which oil droplets or dye injected into the vas deferens moved proximally during electroejaculation in rats and dogs (12, 13). Such retrograde motion likely reflects a rapid intraluminal pressure redistribution that precedes the generation of the ballistic antegrade flow responsible for propelling sperm fluid toward the distal end (1, 9, 13).

### Dynamic sperm flow drives lumen opening in the distal vas deferens

We next investigated how PE-induced contraction alters the structural organization of the vas deferens. To visualize the global architecture of the duct and the distribution of luminal sperm, we performed optical tissue clearing followed by light-sheet microscopy in Pax2-LynVenus mice, which revealed a clearly defined luminal structure filled with densely packed sperm cells at a cm scale (**Figures 3A,S2A**). In the basal state, the luminal structure of epithelial duct exhibits regional variation along the longitudinal axis: it remains open in the proximal region (**Figure 3B**) but progressively narrows and ultimately collapses toward the distal end (**Figure 3C**). Beyond this narrowing point, the distal epithelial sheet exhibits a wrinkled morphology, which we refer to as the distal wrinkle region.

**Figure 3.**
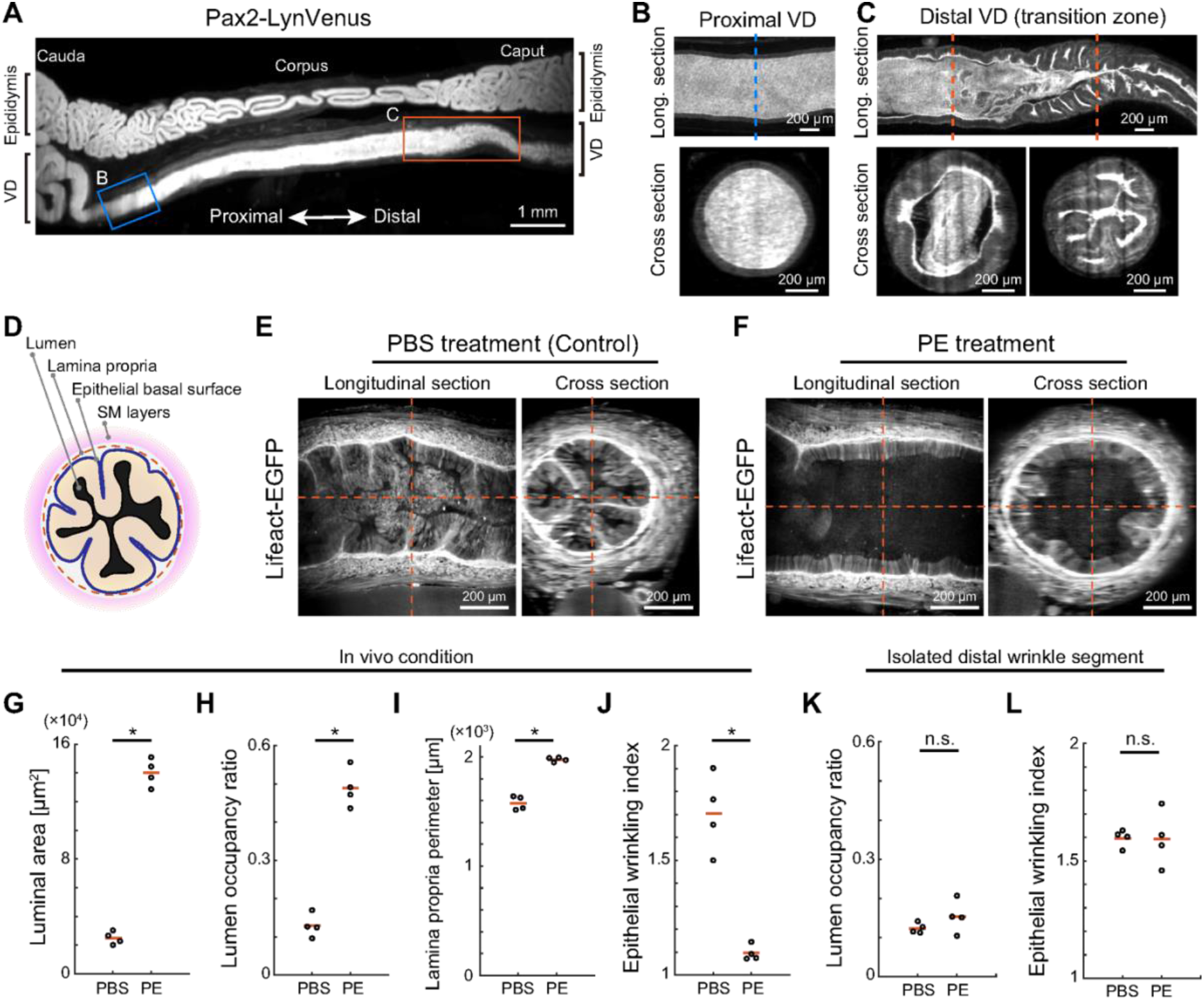
Structural changes in the distal segment of the vas deferens. (A) Light-sheet microscopy image of the vas deferens in a Pax2-LynVenus mouse. The convoluted, wider duct connected to the straight segment at the bottom corresponds to the vas deferens, whereas the folded duct at the top represents the epididymis. Blue and orange boxes indicate the regions shown in panels (B) and (C), respectively. Scale bar, 1 mm. (B,C) Magnified orthogonal views of the proximal (B) and distal (C) regions of the vas deferens. In the distal region, the luminal epithelial surface transitions from smooth to wrinkled, forming a transition zone. Dotted lines in the longitudinal sections indicate the positions of the corresponding cross-sections. Scale bars, 200 µm. (D) Schematic diagram showing the cross-sectional architecture of the distal vas deferens. (E,F) Orthogonal views of the distal wrinkle region in Lifeact-EGFP mice before (E) and after (F) PE treatment. Scale bars, 200 µm. (G–J) Quantification of morphological changes in the distal wrinkle region under in vivo conditions, comparing PBS control and PE-treated samples: (G) luminal area, p=0.0286, (H) lumen occupancy ratio p=0.0286, (I) lamina propria perimeter, p=0.0286, (J) epithelial wrinkling index, p=0.0286. N=4; Mann–Whitney U test. Red bars indicate mean values. (K,L) Quantification of morphological changes in the isolated distal wrinkle segment under ex vivo conditions: (K) lumen occupancy ratio, N=4; Mann–Whitney U test, p=0.343. (L) epithelial wrinkling index, N=5; Mann–Whitney U test, p=0.886. Red bars indicate mean values.

To determine how this distal structure responds to PE stimulation, we used Lifeact-EGFP mice, which enable clear visualization of vas deferens lumen, epithelial layers, lamina propria, and smooth muscle layers (**Figures 3D,S2B**). Tissues were collected and fixed five min after the chemical treatment for whole-tissue imaging. As expected, PBS-treated controls retained a tightly folded epithelial morphology with a collapsed lumen (**Figures 3E,S2C; Movie 9**). In contrast, PE-treated vas deferens displayed an expanded lumen and a smoother epithelial contour with reduced wrinkling (**Figures 3F,S2D; Movie 10**). Quantitatively, PE treatment increased luminal area (**Figure 3G**), elevated lumen occupancy ratio within the epithelial duct (**Figure 3H**), and enlarged perimeter of the lamina propria (**Figure 3I**). To quantify epithelial wrinkles, we defined an epithelial wrinkling index as the ratio of the epithelial basal-surface perimeter to the perimeter of the lamina propria (**Figure 3D**); larger values indicate a more highly wrinkled epithelium. PE treatment significantly reduced this wrinkling index (**Figure 3J**), indicating a morphological transition from a wrinkled to a more flattened epithelial sheet.

To test whether PE itself directly stretches the epithelium, we isolated the distal wrinkle region by cutting at the transition point (**Figure 3C**) and examined PE responsiveness ex vivo. In these isolated segments, PE neither increased luminal occupancy ratio (**Figure 3K**) nor reduced epithelial wrinkling (**Figure 3L**) compared with controls, indicating that PE alone is insufficient to remodel the distal epithelium. We further asked whether activation of a canonical smooth-muscle regulatory pathway is sufficient to remodel the distal epithelium by treating isolated distal wrinkle segments with sodium nitroprusside (SNP), an NO donor that activates soluble guanylyl cyclase–cGMP signaling (30, 31). SNP did not increase lumen area or reduce epithelial wrinkling compared with controls **(Figure S3**), suggesting that SNP-sensitive NO– cGMP signaling alone is insufficient to induce distal epithelial remodeling. Together, these results suggest that distal lumen expansion requires additional tissue-level inputs present in vivo, such as rapid luminal sperm flow and/or mechanical coupling generated by PE-induced contraction of the proximal vas deferens.

### Sperm flow elicits active epithelial remodeling that unfolds the distal vas deferens

To obtain direct evidence that luminal sperm flow drives distal lumen opening, we performed intravital TPEM imaging of the distal wrinkle region in Pax2-LynVenus mice. As expected, PE-induced contraction triggered the previously collapsed lumen to open, with luminal size progressively increasing and epithelial wrinkles unfolding as sperm flowed through the region (**Figure 4A; Movie 11**). Notably, before PE stimulation, sperm were confined within the narrow lumen and remained motionless (**Figures 4B,C left**), even though they displayed robust motility in the proximal vas deferens (**Figure 2B**). Once the lumen opened, however, sperm exhibited directional flow (**Figures 4B,C right**). These observations suggest that, under basal conditions, epithelial wrinkles in the distal region serve as a functional valve that prevents sperm leakage, and that epithelial remodeling occurs in concert with the arrival of luminal sperm flow from the proximal vas deferens.

**Figure 4.**
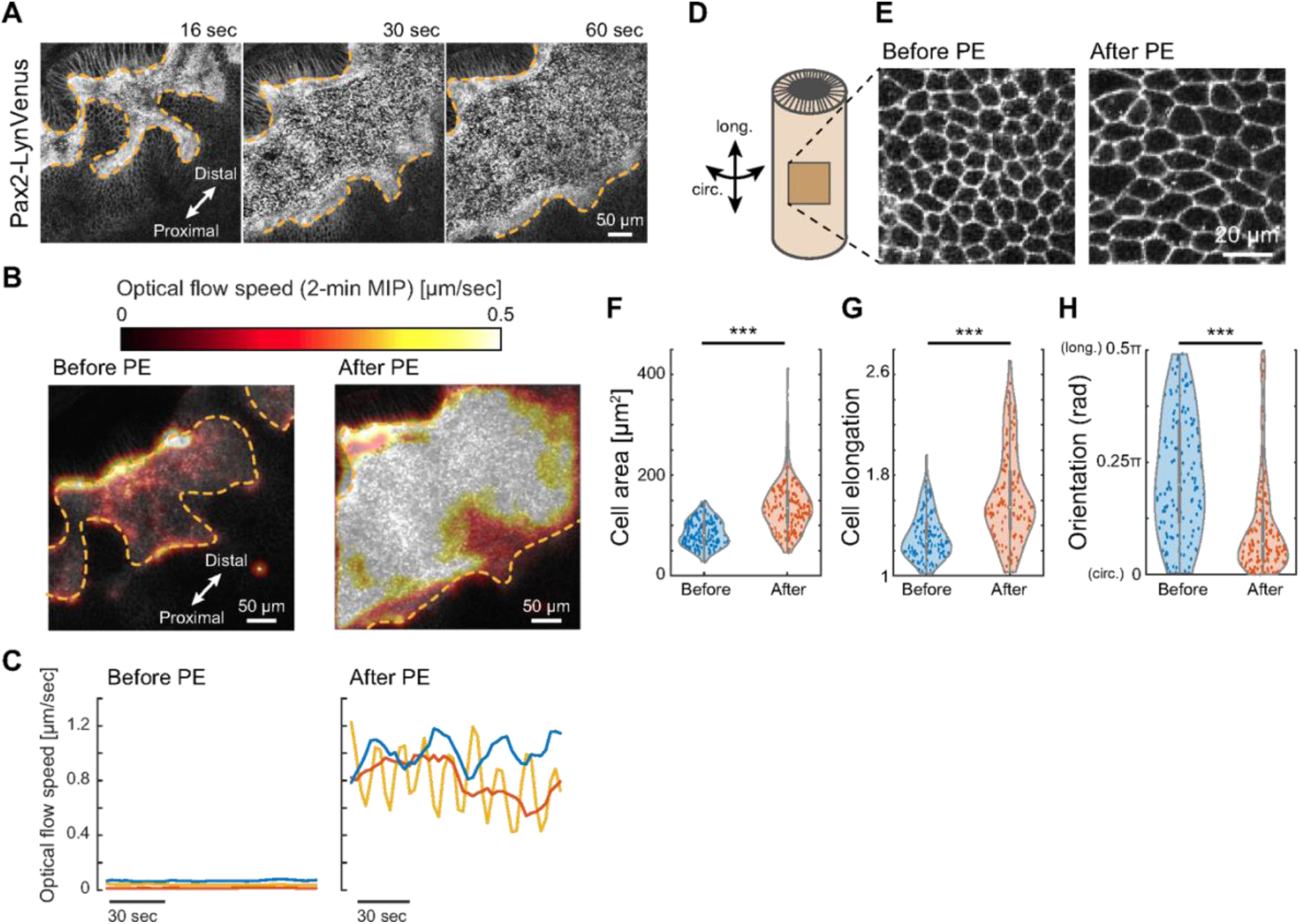
Dynamics of vas deferens epithelium in the distal wrinkle region. (A) Lumen opening in the distal wrinkle region of the Pax2-LynVenus vas deferens following PE treatment. Scale bar, 50 µm. (B) Optical-flow speed maps generated from maximum-intensity projections (MIP) over a 2-min window before and after PE treatment. Orange dotted lines indicate the lumen–epithelium interface. Scale bars, 50 µm. (C) Instantaneous optical-flow speeds before and after PE treatment. N=3. Horizontal time scale bar, 30 sec. (D) Schematic representation of the epithelial duct illustrating two principal orientations: the longitudinal axis and the circumferential axis. (E) Representative epithelial cell morphology before and after PE treatment. Orientation of the longitudinal–circumferential axes follows the scheme in D. Scale bar, 20 µm. (F–H) Quantification of epithelial cell morphological changes in the distal wrinkle region before and after PE treatment: (F) cell area p=4.28×10^-25^, (G) cell elongation (aspect ratio: major axis/minor axis of fitted ellipse), p=2.26×10^-16^, (H) cell orientation (0=circumferential; π/2=longitudinal), p=1.42×10^-14^. N=3, n=150; Mann–Whitney U test.

Since luminal flow imposes shear stress mainly along the longitudinal axis of the duct, the unfolding of longitudinal epithelial wrinkles can be explained as a passive response to fluid shear. However, wrinkles oriented circumferentially also unfolded after PE stimulation (**Figure 3F**), implying additional tissue-level responses beyond passive shear-induced deformation. This prompted us to investigate whether the epithelial tissues that directly face the lumen undergo local strain during luminal flow. By analyzing epithelial cell morphology within the distal wrinkle region, we identified significant changes before and after PE treatment (**Figures 4D,E**). PE stimulation increased epithelial cell area (**Figure 4F**) and enhanced cell elongation specifically along the circumferential axis (**Figures 4G,H**). For both cell area and cell elongation index, PE treatment not only shifted the mean values but also broadened the distributions. This increase in variability is consistent with heterogeneous cell-level responses and suggests that epithelial remodeling is not explained solely by uniform passive unfolding.

### PE stimulation induces distinct spatio-temporal activation patterns of signaling kinases

The preceding experiments showed that PE induces rapid tissue responses on timescales ranging from seconds to minutes, including duct contraction, luminal sperm flow, and distal lumen remodeling. We therefore asked which signaling pathways are activated during these integrated tissue-level events. We selected three candidate kinase pathways representing distinct aspects of vas deferens function: Rho-associated protein kinase (ROCK), because ROKα/ROCK2 is expressed and functionally implicated in murine vas deferens contractility (32, 33); protein kinase A (PKA), because cAMP-associated signaling has been linked to regulation of vas deferens smooth muscle tone (34); and extracellular signal-regulated kinases (ERK), because MAPK/ERK signaling broadly regulates cytoskeletal dynamics, contractile protein expression, and mechanosensitive tissue responses (35–37). To monitor these pathways in vivo, we combined intravital TPEM with Förster resonance energy transfer (FRET)-based kinase biosensors for ROCK, PKA, and ERK (38–40). Because ductal contraction represents the earliest major tissue-level response to PE stimulation, we focused our analysis on smooth muscle cells, both longitudinal and circumferential layers.

We first examined ROCK activity. Before PE stimulation, ROCK activity was low in both longitudinal and circumferential smooth muscle layers. As expected, ROCK activity increased within 10 sec after stimulation and reached a sustained elevated level within 60 sec in both layers (**Figures 5A,A’; Movie 12**). This swift surge in ROCK activity coincided with the onset of vas deferens contraction and is likely to drive both longitudinal shortening and circumferential constriction. Although ROCK activation was detected rapidly in both smooth muscle layers during the early contractile phase, distal lumen expansion occurred subsequently, after luminal sperm flow reached the distal wrinkle region, indicating that circumferential contractile activation and distal valve opening represent temporally distinct components of the PE-induced response.

**Figure 5.**
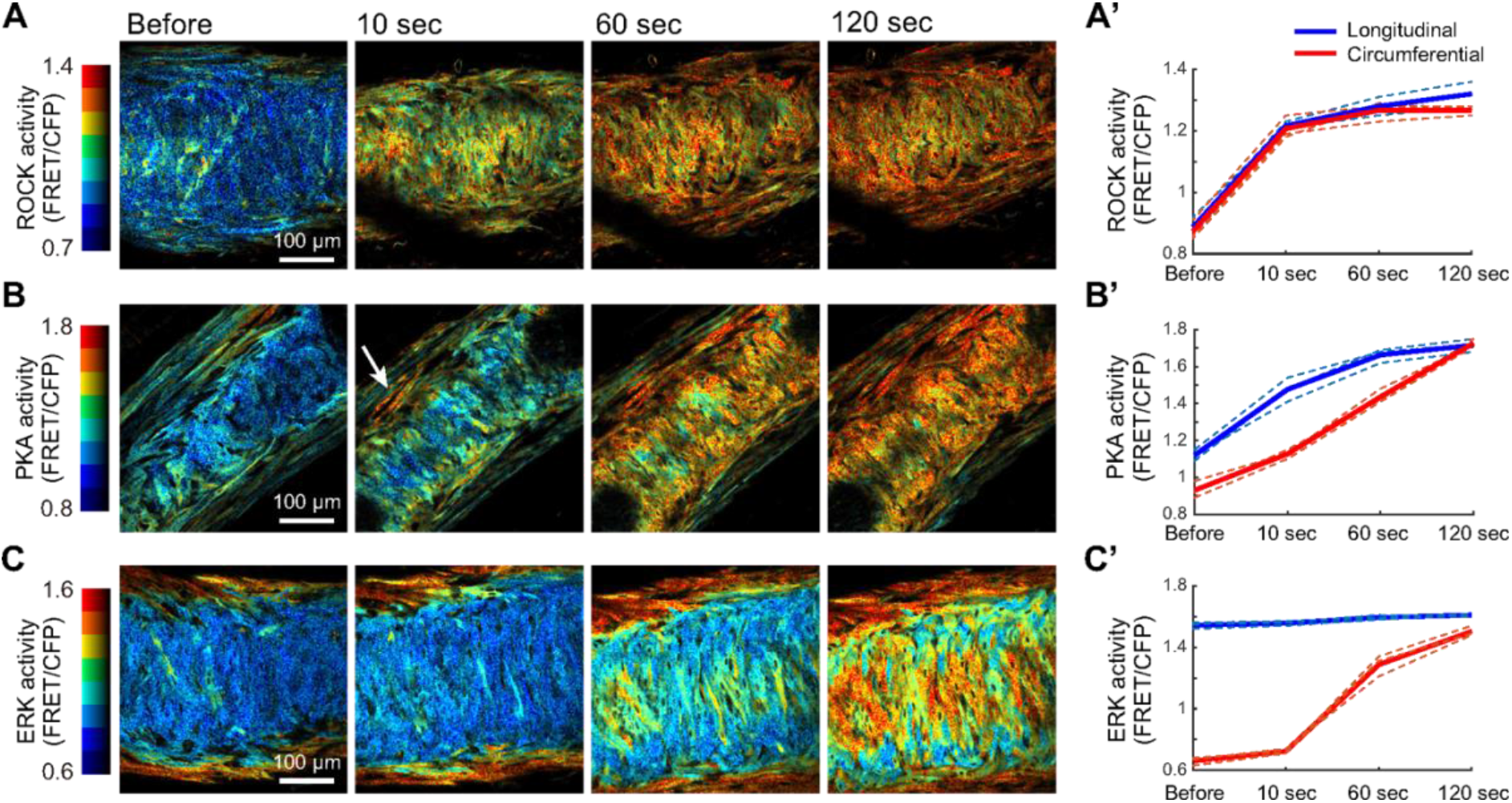
In vivo characterization of kinase activities in the vas deferens. (A–C) Intravital TPEM time-lapse images showing ROCK (A), PKA (B), and ERK (C) activities in the vas deferens before and after PE stimulation. Color denotes kinase activity, and brightness indicates biosensor expression levels. Arrow in (B) highlights longitudinal activation of PKA. Scale bars, 100 µm. (A’–C’) Quantification of ROCK (A’), PKA (B’), and ERK (C’) activities over time. Blue and red traces correspond to longitudinal and circumferential activity, respectively. N=3. Individual samples are shown as dotted lines, with mean values shown as thick solid lines.

Since recent work has implicated cAMP in the regulation of vas deferens smooth muscle (34), we next monitored PKA, also known as cAMP-dependent protein kinase. Under basal conditions, PKA activity was relatively low in both layers, with slightly higher activity in the longitudinal layer than in the circumferential layer (**Figures 5B,B’**). Following PE stimulation, PKA activity in the longitudinal layer increased rapidly, rising by approximately 32% from the basal state within 10 sec (**Figure 5B arrow**), whereas the circumferential layer showed only a 21% increase over the same period. By 120 sec, PKA activity in both layers converged to a similar level, reflecting continued activation in the longitudinal layer and delayed activation in the circumferential layer (**Figures 5B,B’; Movie 13**). Thus, PE evokes a rapid PKA activity increase in longitudinal smooth muscle and a delayed increase in circumferential smooth muscle, suggesting layer-specific roles in shaping vas deferens contractility and luminal sperm transport.

We then evaluated ERK activity, given that the MAPK/ERK pathway broadly regulates contractile protein expression and cytoskeletal dynamics (35–37), although ERK signaling has not previously been examined in the vas deferens. Interestingly, longitudinal smooth muscle exhibited high basal ERK activity that did not increase further upon PE stimulation (**Figures 5C,C’**). In contrast, the circumferential layer showed a clear, time-dependent activation: ERK activity increased by 9% at 10 sec, and continued to rise to 95% and 127% above baseline at 60 and 120 sec, respectively (**Figures 5C,C’; Movie 14**). Accordingly, ERK activity in the distal vas deferens is characterized by sustained, basal signaling in the longitudinal layer and gradual activation in the circumferential layer.

Detectable ERK changes occurred only in the circumferential layer and on a timescale that coincides with the arrival of luminal sperm flow at the distal wrinkle region; thus, we hypothesize that circumferential ERK activation contributes to active lumen remodeling rather than to the initial contractile response. This layer-specific kinase activation pattern suggests a functional specialization in which ROCK drives the rapid global contraction, PKA differentially modulates longitudinal and circumferential tone, and ERK activity in the circumferential layer plays a role in flow-dependent structural remodeling of the distal vas deferens.

### Distinct kinase modules control contraction and tissue flow-dependent remodeling

We further investigated how major signaling kinases influence vas deferens contraction and remodeling by applying pharmacological perturbations to isolated vas deferens tissue. In each experiment, kinase inhibitors or activators were applied for two min, followed by PE stimulation, allowing us to assess how a given kinase affects PE-induced contraction (**Figure 6A**). As a vehicle control, DMSO treatment alone did not alter vas deferens morphology, and subsequent PE stimulation induced 28% contraction (**Figure 6B**). Furthermore, we confirmed that Blebbistatin alone did not cause significant morphological change and its pretreatment abolished PE-induced contraction (**Figure 6C**), corroborating the requirement of actomyosin-based force generation.

**Figure 6.**
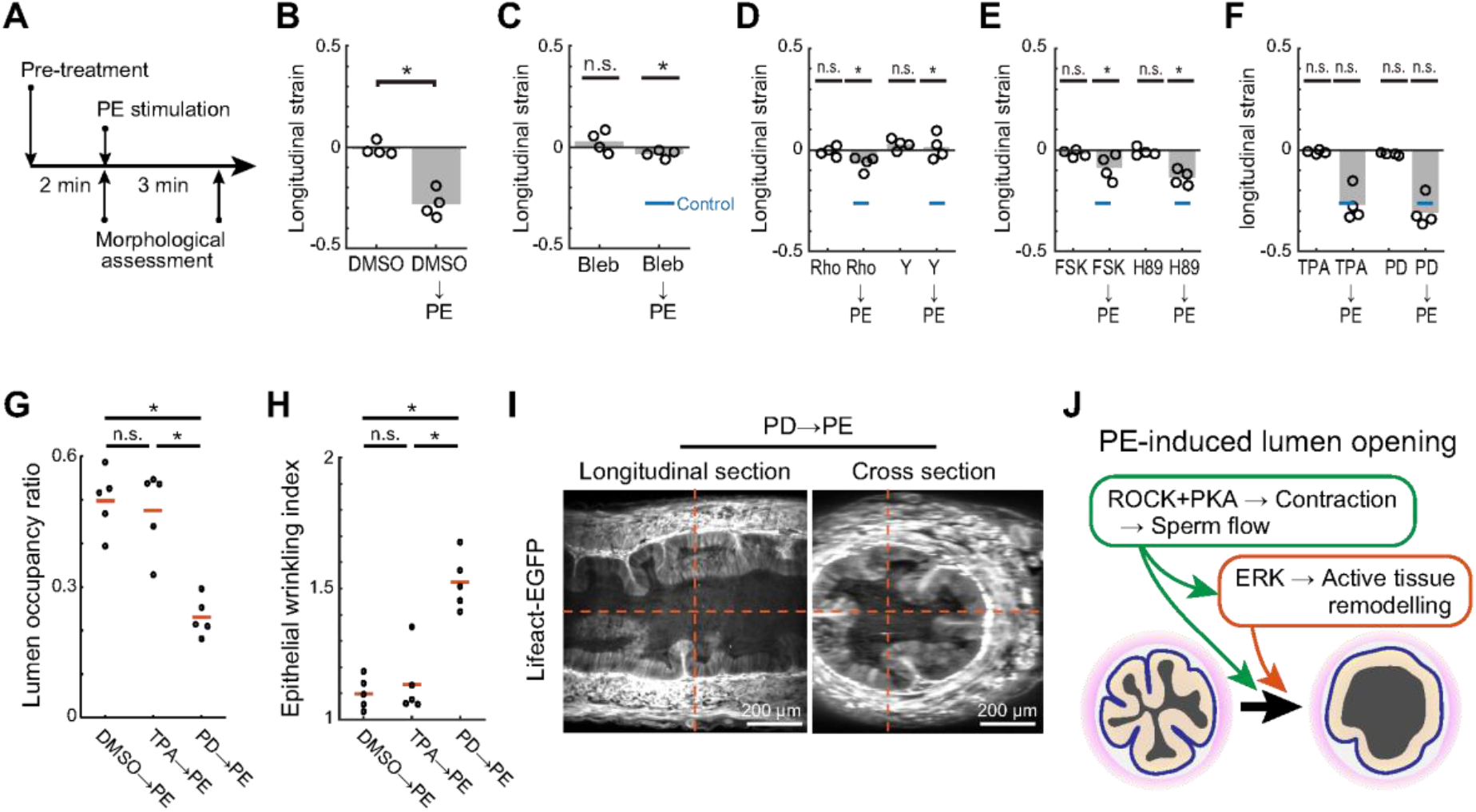
Ex vivo characterization of vas deferens morphological responses to kinase perturbations. (A) Experimental timeline for pharmacological treatment of isolated vas deferens segments. Each segment was pretreated with either an activator or inhibitor, followed by PE stimulation. (B–F) Relative length of vas deferens segments after treatment, normalized to their lengths before treatment. For panels C–F, the pretreatment control was DMSO, and the control for the subsequent PE stimulation was the DMSO→PE condition shown in B. Grey bars indicate mean values. Blue bars in C– F indicate the reference (DMSO→PE) contraction level. N=4 per condition. (B) Calibration of pharmacological treatments. p=0.0286 for DMSO vs. DMSO→PE; Mann–Whitney U test. (C) Contribution of actomyosin contractility assessed using Blebbistatin (100 µM). p=0.486 for Bleb alone and p=0.0286 for Bleb→PE; Mann–Whitney U test. (D) ROCK contribution to longitudinal length changes using Rho activator II (Rho; 10 µg/mL or Y27632 (Y; 30 µM). p=0.886 for Rho, p=0.0286 for Rho→PE, p=0.2 for Y, and p=0.0286 for Y→PE; Mann–Whitney U test. (E) PKA contribution to longitudinal length changes using Forskolin (FSK; 10 µM) or H-89 (H89; 10 µM). p=0.886 for FSK, p=0.0286 for FSK→PE, p=0.686 for H89, and p=0.0286 for H89→PE; Mann–Whitney U test. (F) ERK contribution to longitudinal length changes using 12-O-tetradecanoylphorbol-13-acetate (TPA; 100 nM) or PD0325901 (PD; 1 µM). p=0.343 for TPA, p=1 for TPA→PE, p=0.486 for PD, and p=0.486 for PD→PE; Mann–Whitney U test. (G) Lumen occupancy ratio across drug-treated conditions. Red bars indicate mean values. N=5 per condition; Kruskal-Wallis test, p=0.0092. Tukey–Kramer post-hoc comparisons: p=0.997 for DMSO vs. TPA, p=0.0241 for DMSO vs. PD, and p=0.0197 for TPA vs. PD. (H) Epithelial wrinkling index in the drug treatments. Red bars indicate mean values. N=5 per condition; Kruskal-Wallis test, p=0.009. Tukey–Kramer post-hoc comparisons: p=0.976 for DMSO vs. TPA, p=0.0165 for DMSO vs. PD, and p=0.0294 for TPA vs. PD. (I) Orthogonal views of the distal wrinkle region in Lifeact-EGFP mice treated with PD→PE treatment. Scale bar, 200 µm. (J) Schematic model of PE-induced lumen opening. Contraction-evoked sperm flow passively opens the lumen, followed by ERK-dependent active tissue remodeling that completes lumen opening in the distal vas deferens.

Next, we examined whether acute inhibition or activation of ROCK, PKA, or ERK affects vas deferens morphology. None of these treatments induced significant contraction or relaxation by themselves (**Figures 6D–F**), indicating that short-term perturbation of these kinases does not drive tissue deformation in the absence of PE. Pretreatment with either a ROCK activator (Rho activator) or ROCK inhibitor (Y-27632) blocked the PE-induced contraction (**Figure 6D**), demonstrating that ROCK activation from the basal level is essential for this response. Similarly, pretreatment with Forskolin, which elevates cAMP signaling, or H-89, an inhibitor with known off-target effects, suppressed PE-induced contraction (9% and 14% contraction for Forskolin and H-89 pretreatment, respectively; **Figure 6E**). Together with the PE-induced increase in PKA activity in the longitudinal smooth muscle layer (**Figures 5B,B’**), these findings suggest that cAMP-associated signaling modulates PE-induced vas deferens contraction. In contrast, pretreatment with either an ERK activator (TPA) or a MEK inhibitor (PD0325901, which blocks upstream kinase of ERK) did not prevent PE-induced contraction; PE stimulation elicited contractions comparable to the control condition regardless of prior ERK activation or inhibition (27% and 31% contraction for TPA and PD0325901 pre-treatment, respectively; **Figure 6F**). These results indicate that ERK activity is dispensable for the contractile response.

Given the circumferential ERK activation observed on a slower timescale (**Figures 5C,C’**), we hypothesized that ERK contributes primarily to circumferential lumen remodeling rather than to longitudinal contraction. We therefore examined how ERK signaling regulates the acute morphological response of the lumen in the distal wrinkle region. To test this, we evaluated luminal morphology three min after PE stimulation in tissues pretreated for two min with DMSO, the ERK activator, or the inhibitor. ERK pre-activation did not alter the response to PE; lumen opening and epithelial wrinkle unfolding occurred to a degree comparable to the control (**Figures 6G,H**). In contrast, ERK pre-inhibition impaired both lumen opening and epithelial wrinkle remodeling (**Figures 6G–I**). Compared with PBS treatment alone (**Figures 3G,J**), only partial, rather than complete, lumen opening was observed under PE stimulation following ERK pre-inhibition (**Figures 6G–I**), supporting a sequential, coupled mechanism in the vas deferens: a passive response driven by luminal sperm flow and an active tissue remodeling process triggered by that flow (**Figure 6J**). Together, these findings indicate that flow-associated ERK activation, rather than pre-existing basal ERK activity, drives epithelial wrinkle unfolding.

## DISCUSSION

Here we combined intravital TPEM, light-sheet imaging, and FRET-based kinase biosensing to dissect how luminal sperm transport and vas deferens morphology are coordinated during PE stimulation. Applied here for the first time to the vas deferens, this intravital fluorescence imaging approach revealed the sequence of events that drive ejaculation-like transport under physiologically relevant conditions. We corroborated the intriguing process proposed earlier (7–9) that vas deferens transport involves a rapid retrograde luminal flow, likely reflecting intraluminal pressure redistribution, followed by ballistic antegrade flow that drives sperm from the proximal toward the distal vas deferens. We further showed that this ballistic flow drives opening of the normally collapsed distal lumen and further unfold the epithelial wrinkles. By integrating kinase activity imaging with pharmacological perturbations, we identified distinct signaling modules that control these tissue-level responses: ROCK activity is closely linked to global contraction, cAMP-associated signaling modulates PE-induced contractile responses, and ERK activity is dispensable for the contraction but crucial for circumferential remodeling of the distal epithelium. In parallel, we uncover a previously unexplored role for ERK signaling in supporting smooth luminal sperm flow through flow-dependent lumen opening and epithelial wrinkle unfolding.

Several limitations should be considered when interpreting these pharmacological and signaling data. First, PE stimulation provides a temporally controlled and imaging-compatible trigger of vas deferens contraction, but it does not fully recapitulate physiological neurogenic ejaculation (15). Classical nerve-evoked vas deferens contraction involves sympathetic co-transmission, in which ATP contributes to the rapid twitch component through postjunctional P2X1 receptors, whereas noradrenaline contributes to the slower adrenergic component (16, 41, 42). Thus, the PE-induced response studied here should be interpreted as an ejaculation-like transport model rather than a complete reproduction of endogenous ejaculation. Second, the 10 mM PE concentration used for in vivo imaging was selected empirically to induce reproducible contraction and luminal sperm transport under topical intravital imaging conditions, but it is high relative to receptor-subtype pharmacology and may activate additional adrenergic pathways (26, 43). Therefore, we do not interpret the in vivo response as selective α1-adrenoceptor activation. Third, although α1-adrenoceptor activation is classically coupled to Gq/PLC/IP3–Ca^2+^ signaling, we did not directly dissect this upstream pathway, because broad perturbation of PLC/IP3–Ca^2+^ signaling would be expected to suppress smooth muscle contraction itself (44, 45) and would not readily distinguish contraction initiation from downstream flow-dependent remodeling. Finally, the cAMP/PKA-related data should be interpreted as modulatory rather than causal evidence for contraction. Previous studies have shown that cAMP-elevating treatments, including forskolin, can inhibit vas deferens contraction in rats and guinea pigs (46, 47), consistent with our observation that forskolin suppresses PE-induced shortening in mice. Moreover, forskolin globally elevates cAMP and may act through PKA-dependent and PKA-independent effectors, including EPAC, while H-89 has known kinase off-target effects (48, 49). We therefore interpret these data as evidence that cAMP-associated signaling modulates PE-induced vas deferens responses, rather than as selective evidence that PKA directly drives contraction. Future studies using neurogenic stimulation, receptor-subtype-specific perturbations, and spatially resolved reporters for Ca^2+^, PLC/IP3, PKA, and EPAC will be required to define how endogenous sympathetic signaling engages the tissue-level flow and remodeling dynamics described here.

Based on these findings, we propose that the vas deferens operates as a well-controlled mechanical system that regulates luminal sperm transport through a valve-like distal segment. In the basal state, the distal vas deferens maintains a tightly folded, wrinkled epithelium that collapses the lumen, confines sperm motion, and likely acts as a safeguard against premature leakage. Upon stimulation, PE-induced contraction of the proximal vas deferens generates a rapid antegrade sperm flow that passively pries open the distal lumen, initiating the first, flow-driven phase of wrinkle unfolding. This is followed by an active remodeling phase in which vas deferens tissues, through a flow-sensing mechanism, engage ERK signaling in circumferential smooth muscle, and potentially epithelium, to stabilize lumen opening and complete wrinkle unfolding. Thus, beyond the well-established insights from electrophysiology and genetic models (15, 17–19), our study adds a mechano-signaling perspective, clarifying how coordinated ROCK–PKA–ERK activity and tissue mechanoresponsiveness together ensure robust, directional, and efficient luminal sperm transport. More broadly, these findings provide insight into how valve-like structures in the reproductive tract, such as the Sertoli valve and the uterotubal junction, are organized and regulated (50–53).

This study proposes a new view of vas deferens behavior during the ejaculation process and, at the same time, highlights several open questions. The first key issue concerns mechanosensing of luminal flow. How is the hydrodynamic input generated by luminal sperm fluid detected, and which cellular machinery implements this sensing? A plausible candidate structure is the stereocilia on epithelial cells that directly face the lumen. In other systems, such as the cochlear duct, actin-based stereocilia function as mechanosensory organelles, where fluid-driven bundle deflection opens ion channels and triggers Ca^2+^ influx (54, 55). Vas deferens epithelial cells in both rodents and humans possess similar apical specializations (7), yet their mechanosensory roles remain essentially unexplored. It will be important to determine whether these stereocilia participate in sensing luminal flow and in initiating the ERK-dependent signaling (56) that we associate with wrinkle unfolding and lumen remodeling. The second set of questions relates to inter-tissue crosstalk during mechanotransduction. Although the primary flow-sensing interface likely resides in the epithelium, the ultimate control of duct geometry also lies in the surrounding smooth muscle layers. Signals originating in epithelial cells whether chemical, mechanical, or both, must therefore be transmitted across the lamina propria, a matrix-rich compartment containing basement membrane and associated cells, to coordinate smooth muscle behavior. How these signals traverse the lamina propria, and how geometric consistency between the epithelial layer, lamina propria, and smooth muscle is maintained during acute morphogenetic changes, remain unclear. Finally, the detailed cellular mechanisms of tissue remodeling are still to be defined: does lumen expansion primarily reflect circumferentially biased stretching of epithelial sheets through cell softening, or does it involve active cell rearrangement and oriented cell elongation in the circumferential direction? Addressing these questions will require volumetric fast live imaging, mechanical perturbations, and targeted manipulation of candidate mechanosensors and signaling pathways (1, 57). Building on the mechano-signaling framework established here, we anticipate that such work will not only deepen our understanding of male reproductive physiology but also provide general principles for how tubular organs regulate material transport through flow-dependent tissue remodeling.

## MATERIALS AND METHODS

### Animals

All animal experiments were approved by the local institutional animal ethics committees (R22-0765, BR23-1140, and MedKyo 21043) and were conducted in accordance with institutional and national guidelines for the care and use of laboratory animals. Adult male C57BL/6J mice were purchased from InVivos Pte Ltd and adult male B6N-Tyrc-Brd/BrdCrCrl mice were purchased from Charles River Laboratories Japan, Inc. Additional transgenic mouse strains were obtained from in-house breeding colonies and were originally generated or provided as follows: Pax2-LynVenus mice were generated and reported previously (28, 29); Pih1d3 knockout mice were reported elsewhere (27) and were obtained from RIKEN BioResource Research Center through the National Bio-Resource Project of the MEXT, Japan (RBRC04704); Lifeact-EGFP mice were reported previously (58) and were provided by Dr. Takashi Hiiragi (EMBL Heidelberg, Germany); ROCK FRET biosensor mice were described previously (38); PKAchu mice were described previously (39, 59) and were obtained from the National Institutes of Biomedical Innovation, Health and Nutrition (NIBIOHN), Japan (nbio185); hyBRET-ERK-NES mice were described previously (40, 60) and were obtained from NIBIOHN (nbio325). All mice were housed under specific pathogen–free conditions in an AAALAC-accredited facility on a 12-h light/dark cycle with ad libitum access to food and water. Age-matched adult males (8–20 weeks) were used for all experiments unless otherwise indicated.

### Small molecules

The following small molecules were used: Forskolin (Cayman Chemical, #11018), H-89 (Cayman Chemical, #10010556), Rho Activator II (Cytoskeleton, #CN03), Sodium nitroprusside dihydrate (Sigma-Aldrich, #71778), Y-27632 (FUJIFILM Wako Pure Chemical Corporation, #259-00613), 12-O-tetradecanoylphorbol-13-acetate (LC Laboratories, #P-1680), PD0325901 (FUJIFILM Wako Pure Chemical Corporation, #62-25291), (-)-Blebbistatin (FUJIFILM Wako Pure Chemical Corporation, #027-17043), (R)-(-)-Phenylephrine hydrochloride (FUJIFILM Wako Pure Chemical Corporation, #163-11791),

### Stereomicroscopy

Stereomicroscope imaging was performed using either an SZX16 stereomicroscope (Evident) equipped with a DP80 color CCD camera (Olympus/Evident) or a DP28 color CMOS camera (Evident), or an SZ61 stereomicroscope (Evident) equipped with an SS500MC color sCMOS camera (Meiji Techno Co., Ltd.). Images were acquired under bright-field illumination using the corresponding camera acquisition software.

### Intravital two-photon microscopy

Adult male mice were anesthetized by induction in an enclosed chamber with 4–5% vaporized isoflurane in oxygen, followed by maintenance at 1–3% isoflurane delivered through a nose cone at a flow rate of 1–2 L/min. Anesthetized animals were positioned on a temperature-controlled heated stage maintained at 37 °C, using a customized metal plate with a 40 × 21 mm opening (Tokai Hit) to accommodate the exposed vas deferens. The isoflurane concentration was adjusted as needed to maintain a stable surgical plane of anesthesia, assessed by the pedal reflex and the absence of voluntary movement. For surgical exposure of the vas deferens, a midline scrotal incision was made using fine surgical scissors without the need for fur removal. The testis and epididymal structures were gently exteriorized to expose the vas deferens, and all tissues were kept hydrated with sterile phosphate-buffered saline (PBS). The mouse was positioned prone on the heated stage such that the vas deferens rested against a #1 coverslip mounted over the microscope objective, enabling inverted imaging without additional physical restraint. Two-photon imaging was performed on inverted multiphoton microscopes (AX R MP, Nikon; FV1200MPE-IX83, Evident) equipped with either a 25× silicone-immersion objective (NA=1.05, WD=0.55 mm, MRD73250, Nikon) or a 30× silicone-immersion objective (NA=1.05, WD=0.8 mm, UPLSAPO30XS, Evident). Excitation wavelengths were set to 840 nm for the CFP channel of FRET biosensors, 930 nm for Lifeact-EGFP, and 945 nm for Pax2-LynVenus (InSight DeepSee, Spectra-Physics). For intravital kinase imaging, we used transgenic mouse strains expressing FRET biosensors reporting the activities of ROCK, PKA, or ERK. These biosensors comprise a CFP donor and YFP acceptor connected by a kinase-specific substrate peptide and a phospho-binding domain, allowing real-time readout of kinase activity based on changes in the FRET/CFP emission ratio. For FRET image analysis, images were denoised using a 3×3 median filter, and background fluorescence was subtracted separately from the FRET and CFP channels. FRET/CFP ratio images were generated using a custom MATLAB (MathWorks) script. In the resulting pseudocolor maps, hue represents the FRET/CFP ratio and brightness corresponds to the FRET-channel intensity.

### Optical tissue clearing and light-sheet microscopy

Whole vas deferens tissues were optically cleared using CUBIC-L/R reagents (61) and imaged by light-sheet fluorescence microscopy. Dissected vas deferens tissues were fixed in 4% paraformaldehyde (PFA) in PBS overnight at 4°C. The following day, samples were washed three times in PBS for 10 min each at 23°C and incubated in 50% (v/v) CUBIC-L diluted in PBS for 6 h, followed by incubation in 100% CUBIC-L with gentle shaking at 37°C overnight. The CUBIC-L solution was refreshed on the next day, and samples were further incubated for an additional 24 h at 37°C. After clearing with CUBIC-L, tissues were washed in PBS three times for 1 h each at 23 °C and transferred to 50% (v/v) CUBIC-R in PBS for 6 h, followed by 100% CUBIC-R with shaking at 37°C until the samples became optically transparent.

Whole-organ fluorescence imaging was performed using a custom-built light-sheet fluorescence microscopy system constructed around an MVX10 macro zoom microscope system (Evident). The optical configuration of the custom system is described in the legend of **Figure S3A**. Briefly, the system was equipped with an OBIS 488 nm laser (Coherent) for fluorescence excitation. The illumination path consisted of collimation optics, a galvo scanning mirror, relay optics, and cylindrical lenses. The galvo mirror rapidly swept the excitation beam across the specimen, while the cylindrical optics shaped the beam to generate a scanned light sheet at the specimen plane, following the principle of digitally scanned light-sheet fluorescence microscopy (62). Fluorescence emission was collected through the MVX10 detection path and recorded using an sCMOS camera (Neo, Andor). Images were acquired using either a 0.63× objective lens with a waterproof cover (NA=0.15, WD=30 mm; Evident) or a 1× objective lens with a waterproof cover (NA=0.25, WD=10 mm; Evident). The 0.63× objective lens was used for large-scale anatomical imaging, whereas the 1× objective lens was used for higher-magnification imaging and wrinkle analysis. For refractive-index matching during imaging, tissues were immersed in a 1:1 mixture of silicone oil TSF4300 (Momentive Performance Materials; RI=1.498) and mineral oil (Sigma-Aldrich; RI=1.467), following (61).

### Optical flow analysis

Optical flow analysis was performed in MATLAB (MathWorks) using the Farnebäck dense-flow algorithm to quantify pixel-wise motion of luminal sperm and tissue structures. Time-lapse image sequences were first registered to correct for sample drift. Dense optical flow fields were then computed for each pair of consecutive frames using the opticalFlowFarneback function from the Computer Vision Toolbox, with parameter settings optimized for the spatial scale and velocity range characteristic of vas deferens dynamics. The resulting optical flow vectors were visualized either as instantaneous velocity arrows or as color-coded speed maps. All analyses and visualizations were performed using custom MATLAB scripts.

### Quantification of directional order parameter

The directional order parameter was used to quantify the degree of collective alignment in sperm motion. It was defined as 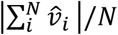, where 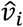 denotes the unit velocity vector within the i-th sub-window, and *N* is the total number of sub-windows. Velocity vectors were obtained from optical flow analysis of time-lapse image sequences. The magnitude of directional order parameter ranges from 0, indicating completely disordered motion, to 1, indicating perfectly aligned collective motion.

### Quantification of cross-sectional morphological parameters

For quantitative analysis, images acquired from either Lifeact-EGFP or Pax2-LynVenus reporter mice were used. Three regions of interest were manually delineated in Fiji/ImageJ: (1) the lumen, (2) the epithelial basal surface, defined as the interface between the basal side of the epithelium and the lamina propria, and (3) the convex hull of the lamina propria. The convex hull was defined as the smallest convex boundary enclosing the lamina propria region, based on intensity and texture profiles identified by visual inspection. For each region, area and perimeter were measured using the ImageJ built-in measurement function. The lumen occupancy ratio was calculated as the ratio of the lumen area to the area of the convex hull of the lamina propria. The epithelial wrinkling index was defined as the ratio of the perimeter of the epithelial basal surface to that of the convex hull of the lamina propria. An index value of 1 indicates a smooth, non-wrinkled basal surface, representing the minimum value, whereas higher values reflect increasing degrees of epithelial wrinkling.

### Statistical hypothesis testing

The number of cells analyzed (n) and the number of biological replicates (N) are reported in the figure legends. No statistical methods were used to predetermine sample sizes, and no randomization was performed. Statistical tests, sample sizes, and corresponding p-values are described in the figure legends. We considered p<0.05 statistically significant for two-tailed tests, unless otherwise stated. Significance levels are denoted as follows: * (p<0.05), ** (p<0.01), *** (p<0.001), and n.s. (not significant; p≥0.05).

### Software and data visualization

Digital image processing, quantitative analysis, and visualization were conducted using MATLAB R2019b, R2022b, and R2024b (MathWorks) and ImageJ (National Institutes of Health). Truncated violin plots were generated using a MATLAB function sourced from MATLAB File Exchange (63).

## ACKNOWLEDGEMENTS

This work was supported by Singapore Ministry of Education (MOE) Academic Research Fund (AcRF) Tier 2 (MOE-T2EP30223–0010), the National Research Foundation, Singapore (NRF) under its Mid-sized Grant (NRF-MSG-2023–0001), and Hakubi fellowship grant of Kyoto University. Light-sheet microscopy was supported by the Kyoto University Live Imaging Center, and 3D image processing was supported by Hui Ting Ong from the Microscopy Core at the Mechanobiology Institute. We thank all members of the laboratory for their helpful discussions.

## AUTHOR CONTRIBUTIONS

Conceptualization, GY, TH

Methodology, GY, TH

Investigation, NNSY, QYL, TH

Data curation, NNSY, QYL, TH

Formal analysis, NNSY, TH

Writing – original draft, NNSY, QYL, TH

Writing – review & editing, GY, TH

Supervision, TH

Funding acquisition, TH

Resources, TH

Visualization, NNSY, TH

## DATA AVAILABILITY

All data supporting the findings of this study are provided in the article and Supplementary Materials. Source data underlying the quantitative plots are provided as Dataset S1.

